# The dynamic nature of neurotensin receptor 1 (NTS_1_) allostery and signaling bias

**DOI:** 10.1101/2022.11.25.517797

**Authors:** Fabian Bumbak, Asuka Inoue, Miquel Pons, Juan Carlos Paniagua, Fei Yan, Hongwei Wu, Scott A. Robson, Ross A. D. Bathgate, Daniel J. Scott, Paul R. Gooley, Joshua J. Ziarek

## Abstract

The neurotensin receptor 1 (NTS_1_) is a G protein-coupled receptor (GPCR) with promise as a drug target for the treatment of pain, schizophrenia, obesity, addiction, and various cancers. A detailed picture of the NTS_1_ structural landscape has been established by X-ray crystallography and cryo-EM and yet, the molecular determinants for why a receptor couples to G protein versus arrestin transducers remain poorly defined. We used ^13^C^ε^H_3_-methionine NMR spectroscopy to show that phosphatidylinositol-4,5-bisphosphate (PIP2) promotes transducer complexation not by dramatically altering the receptor structure but by strengthening long-range allosteric connections, in the form of correlated conformational kinetics, between the orthosteric pocket and highly-conserved activation motifs. β-arrestin-1 further remodels the receptor ensemble by reducing conformational exchange kinetics for a subset of resonances, whereas G protein coupling has little to no effect on the rate. A β-arrestin biased allosteric modulator transforms the NTS_1_:G protein complex into a concatenation of substates, without triggering transducer dissociation, suggesting that it may function by stabilizing signaling incompetent G protein conformations such as the non-canonical state. Together, our work demonstrates the importance of kinetic information to a complete picture of the GPCR activation landscape.

## INTRODUCTION

Cells rely on membrane-embedded receptors to maintain awareness of the extracellular environment without compromising membrane integrity. The G protein-coupled receptor (GPCR) superfamily is the largest group among such eukaryotic cell surface receptors, comprising more than 800 proteins^1,2^. They are ubiquitously expressed throughout the human body and are pivotal in a broad range of physiological processes including vision, taste, sense of smell, nervous functions, immune regulation, reproduction, and cancer^3,4^. Ligand binding at the extracellular, orthosteric site allosterically induces conformational changes across the GPCRs’ signature seven transmembrane (7TM) helices that prime the intracellular face for interaction with transducer proteins such as G proteins, β-arrestins (βArr), and G protein-coupled receptor kinases (GRKs)^5^. The neurotensin receptor 1 (NTS_1_) is a high-affinity target for the endogenous 13-residue peptide agonist neurotensin (NT)^6^. NT functions as a neuromodulator of the central nervous system (CNS) as well as a paracrine and endocrine modulator of the digestive tract and cardiovascular system^7^.

The strong expression overlaps of the dopamine system with both NT and NTS_1_ has led to considerable evidence for functional synergy in psychostimulant and opioid drug addiction^8-12^. Despite the long-standing interest in NTS_1_ as a potential therapeutic target for substance use disorders (SUDs), the handful of small-molecule NTS_1_ agonists and antagonists that have been developed all suffer from on-target side effects such as hypothermia^13,14^, hypotension^15^, and impaired motor control^15,16^. The classical model of GPCR activation implies that ligand-bound receptors signal equally (aka balanced) through G protein and β-arrestin (βArr) transducer pathways. The recent recognition of biased signaling, in which ligands preferentially activate one transducer pathway over the other, offers a new treatment avenue that may reduce on-target side effects^17-19^. A high-throughput functional screen and ligand optimization campaign targeting NTS_1_ led to the development of ML314, an allosteric ligand that selectively activates βArr2 pathways without stimulating the Gq pathways (i.e. βArr2-biased allosteric modulator; BAM), which reduces addictive behaviors toward methamphetamine and cocaine in several mouse models^20,21^.

Yet, the molecular determinants for why a ligand promotes G protein:receptor versus arrestin:receptor complexation remain poorly defined. The predominant hypothesis is that agonists stabilize distinct receptor conformations to preferentially activate one pathway over another. Recent cryo-EM structures of NTS_1_:βArr1 and NTS_1_:G protein, however, reveal a remarkably conserved receptor architecture with a 0.67 Å all-atom RMSD^22-25^. Solution NMR has proven indispensable for identifying receptor conformers that are invisible to static structural methods^26^, but few studies have investigated G protein^27-29^ or βArr^30-32^ ternary complexes; to date, the only GPCRs characterized by NMR in complex with mimetics of both transducers are NTS_1_^32^ and the β2-adrenergic receptor^30,31,33,34^. Here, we uniformly-label ^13^C^ε^H_3_-methionine residues located within the NTS_1_ transmembrane bundle and near the ligand-binding site to demonstrate how ligands and PIP2 dynamically prepare the receptor for transducer interaction. The differential conformational kinetics upon coupling to βArr and G protein transducer molecules suggests a role for dynamics in functional selectivity.

## RESULTS

### PIP2 strengthens correlated motions of the orthosteric pocket and PIF motif

Phosphatidylinositol-4,5-bisphosphate (PIP2), and its analog (C8-PIP2; here termed PIP2), have been shown to generally enhance the ability of GPCRs to activate G proteins^35,36^ and maximally-recruit arrestin^22^. We used a previously characterized minimal methionine enNTS_1_ variant (herein enNTS_1_ΔM4) to explore the molecular mechanism of PIP2’s positive allosteric modulation^37^. As illustrated in Figure 1A, enNTS_1_ΔM4 retains six endogenous methionine residues (M204^4.60^, M208^4.64^, M244^5.45^, M250^5.51^, M330^6.57^ and M352^7.36^) (superscript refers to Ballesteros-Weinstein numbering^38^). Two-dimensional (2D) ^1^H-^13^C heteronuclear multiple quantum correlation (HMQC) spectra were collected for apo, NT8-13 bound, ML314 bound, and NT8-13:ML314 bound [^13^C^ε^H_3_-methionine]-enNTS_1_ΔM4 in the presence and absence of PIP2 (Figure 1B-E). All NMR spectra were collected at 65 μM [^13^C^ε^H_3_-methionine]-enNTS_1_ΔM4 with identical acquisition, processing, and display parameters; thus, we can directly compare both the chemical shift values (i.e. structure) and signal intensities (i.e. dynamics) for each liganded state.

**Figure 1.**
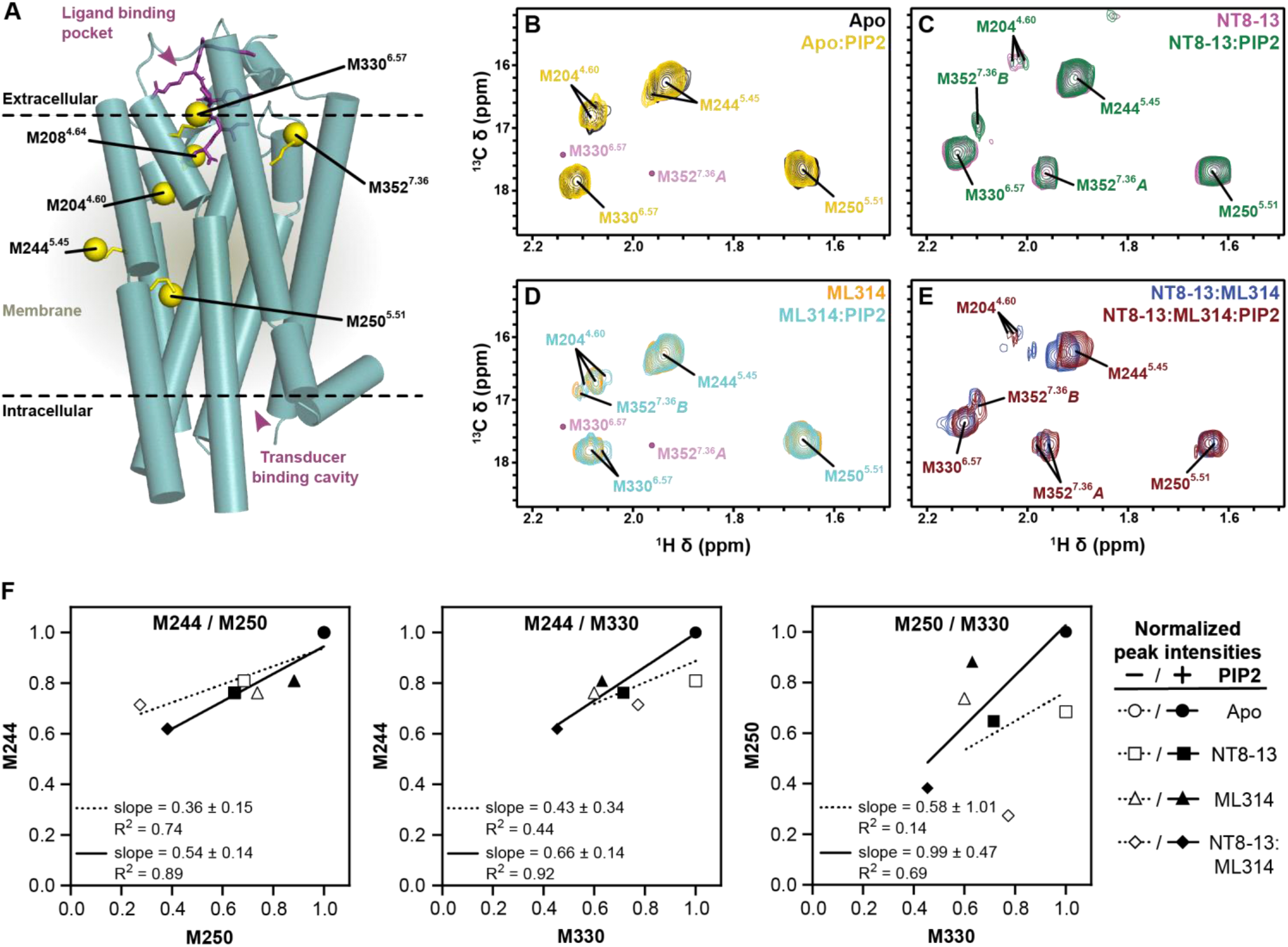
Effect of PIP2 on enNTS_1_ΔM4 ^13^C^ε^H_3_-methionine chemical shifts. A) Cylindrical representation of thermostabilized rNTS_1_ (PDB 4BWB) with labelled methionine methyl groups shown as yellow spheres (superscript - Ballesteros-Weinstein nomenclature^38^) and NT8-13 shown as purple sticks. Overlays of Apo-state (B), NT8-13 (C), ML314 (D), and NT8-13 & ML314 (E) bound ^1^H-^13^C HMQC spectra in the absence and presence of 130 μM (2x molecular equivalents over enNTS_1_) PIP2. The corresponding peak intensities are plotted in Supplemental Figure S1. Pink dots (panels B and D) indicate NT8-13:enNTS_1_ΔM4 M330^6.57^ and M352^7.36^*A* peak positions from panel C. All spectra were recorded at 600 MHz, in 3 mm thin wall precision NMR tubes (Wilmad), with enNTS_1_ΔM4 concentrations of 66 μM. F) Pairwise correlation plots of normalized peak intensities (integrated peak volumes) for M244^5.45^, M250^5.51^, and M330^6.37^ resonances. Symbols correspond to Apo (circle), NT8-13-(square), ML314-(triangle) and NT8-13 & ML314-bound (diamond) enNTS_1_ΔM4 in the presence (filled symbol; solid line) and absence (empty symbol; dotted line) of PIP2. The corresponding slopes and R^2^ values are indicated within each panel.

Except for the NT8-13:ML314 bound state, PIP2 induced only subtle [^13^C^ε^H_3_-methionine]-enNTS_1_ΔM4 chemical shift perturbations indicating minimal effect on the overall structure (Figure 1B-E). At the same time, PIP2 distinctly tunes methionine peak intensities of each receptor complex compared to the ligands alone (Figure S1). Intensity decreases are usually caused by line broadening that may signify either changes in the intrinsic transverse relaxation rate (R_2_) and/or exchange broadening. R_2_ relaxation results from physical properties of the methyl group on the pico-nanosecond (ps-ns) timescale, while exchange broadening reflects conformational interconversion on the micro-millisecond (μs-ms) timescale^39^. In the apo state, PIP2 rigidifies M330^6.57^ adjacent to the ligand-binding pocket even as it increases the μs-ms dynamics of M204^4.60^ at the base of the pocket and M250^5.51^ of the connector region (Figures 1B and S1). PIP2 universally increases μs-ms dynamics across the NT8-13 bound receptor (Figures 1C and S1) whereas it rigidifies the ML314:enNTS_1_ΔM4 complex throughout the extracellular region (M330^6.57^), near the base of the orthosteric pocket (M204^4.60^), and surrounding the PIF motif (M2445.45 and M2505.51; Figures 1D and S1). When both agonist and BAM are present, PIP2 perturbs the M204^4.60^, M244^5.45^, and M250^5.51^ chemical shifts (i.e. pushes the structural equilibrium) towards the NT8-13 bound state (Figure 1E). At the same time, PIP2 selectively stabilizes only the upfield resonance of M330^6.57^ which is a doublet in the NT8-13:ML314:enNTS_1_ΔM4 spectra (Figure 1E). Taken together, these chemical shift changes may reflect PIP2’s balanced potentiation of NTS_1_:transducer coupling and activation^22,35,36^.

The M352^7.36^ peak pattern in both NT8-13:enNTS_1_ΔM4 (Figure 1C) and NT8-13:ML314:enNTS_1_ΔM4 (Figure 1E) complexes reflects a multi-state equilibrium with mixed exchange regimes^40^. At first approximation, the presence of three peaks (states A, B, and C) indicates that M352^7.36^ exchanges between three chemical environments qualitatively on the slow (i.e. ms-s) timescale^37,41^. A previous density functional theory-guided NMR analysis^37^ suggests that state A represents tight packing of TM1/6/7 and lid-like engagement of the N-terminus against the bound NT8-13 as observed in the NTSR1-H4_X_:NT8-13 X-ray structure (PDB 6YVR^42^), whereas M352^7.36^*B* reflects detachment of the receptor N-terminus and local stabilization of extracellular TM1/6/7 as observed in the NTSR1-H4_X_:SR142948A structure (PDB 6Z4Q^42^). In the agonist bound state, PIP2 modestly reduced the total (sum of states A and B) M352^7.36^ peak intensity by 22.6%, which suggests a subtle adjustment towards a faster state A ⇔ state B interconversion rate although still within the ms-s timescale^41^. At the same time, the relative M352^7.36^*B* population increased by 138.8% while the M352^7.36^*A* state was effectively unchanged (Figures 1C and S1). One potential explanation for this behavior is that state B is a composite of two microstates exchanging on the intermediate-fast (μs-ms) timescale. When both agonist and BAM are present, PIP2 increases the total (sum of states A and B) M352^7.36^ peak intensity by 164% without changing the relative state A and B populations (66% and 34%, respectively), which would correspond to a reduction of the A ⇔ B interconversion rate on the slow (ms-s) timescale^41^. The M352^7.36^*B* resonance is simultaneously perturbed upfield in the ^1^H dimension and downfield in the ^13^C dimension (Figure 1E). There are several possible explanations for this behavior: i) a change in the relative populations of fast-exchanging (μs-ms) microstates that comprise state B, ii) structural changes of the M352^7.36^*B* chemical environment itself, and iii) remodeling of both the thermodynamic and kinetic properties of states A and B.

To further explore PIP2-mediated cooperativity between the orthosteric pocket (M330^6.57^) and connector region (M244^5.45^ and M250^5.51^), we measured the pairwise correlation of normalized peak intensities for each liganded state (Figure 1F). This analysis relies on the assumption that residues involved in the same allosteric network will exhibit a concerted response – reminiscent of chemical shift covariance analysis (CHESCA)^43^ and methionine chemical shift-based order parameter analysis^37,44^. Residues M244^5.45^ and M250^5.51^ which are located before and after P^5.50^, respectively, inform on the dynamics across the TM5 kink; the effect is relatively consistent regardless of PIP2, although the better linear correlation suggests an improved dynamic scaling between those residues. Similar results are observed for the pairwise correlation of M244^5.45^ and M330^6.57^, hinting at a subtle allosteric effect throughout the extracellular vestibule. Lastly, we looked at the correlation between M250^5.51^ and M330^6.57^. In the absence of PIP2, the two residues are effectively uncoupled (R^2^ = 0.14). Addition of PIP2 increases the R^2^ to 0.69 and the slope to 0.99, indicating a clear allosteric coupling between the orthosteric pocket and connector region that may provide a mechanism for PIP2’s ability to stabilize active states^35,36^. For the remainder of this study, unless otherwise stated, all samples include PIP2.

### βArr1 alters the kinetic landscape of the NTS_1_ conformational ensemble

Recent cryo-EM structures of the NTS_1_:βArr1 complex required either protein fusion^24^ or intermolecular cross-linking^22^ to stabilize intrinsic dynamics, suggesting that NMR could provide additional information on the nature of these underlying motions. We utilized the pre-activated βArr1-3A variant to maximize the affinity for unphosphorylated enNTS_1_ΔM4^45,46^. enNTS_1_ΔM4:βArr1-3A ternary complex formation resulted in the appearance of two additional methionine peaks (Figure 2A; asterisks). Collecting a ^1^H-^13^C HMQC spectrum using unlabeled enNTS_1_ΔM4, we show that both peaks belong to βArr1-3A; while the major resonance is always detectable, the minor peak is only visible in the presence of the receptor (Figure 2B). Both resonances are unobservable when the experiment is repeated using βArr1 C-terminally truncated after N382 (βArr1-ΔCT^46^). As arrestin recruitment requires displacement of its self-associated C-tail^24,47^, we conclude that the major and minor resonances correspond to βArr1’s C-terminal M411 in the bound-basal and dissociated receptor-bound (or post-receptor-bound) state, respectively. In the absence of PIP2, βArr1-3A promotes limited [^13^C^ε^H_3_-methionine]-enNTS_1_ΔM4 spectral changes further supporting the lipid’s role in high affinity transducer complexation (Figure S2).

**Figure 2.**
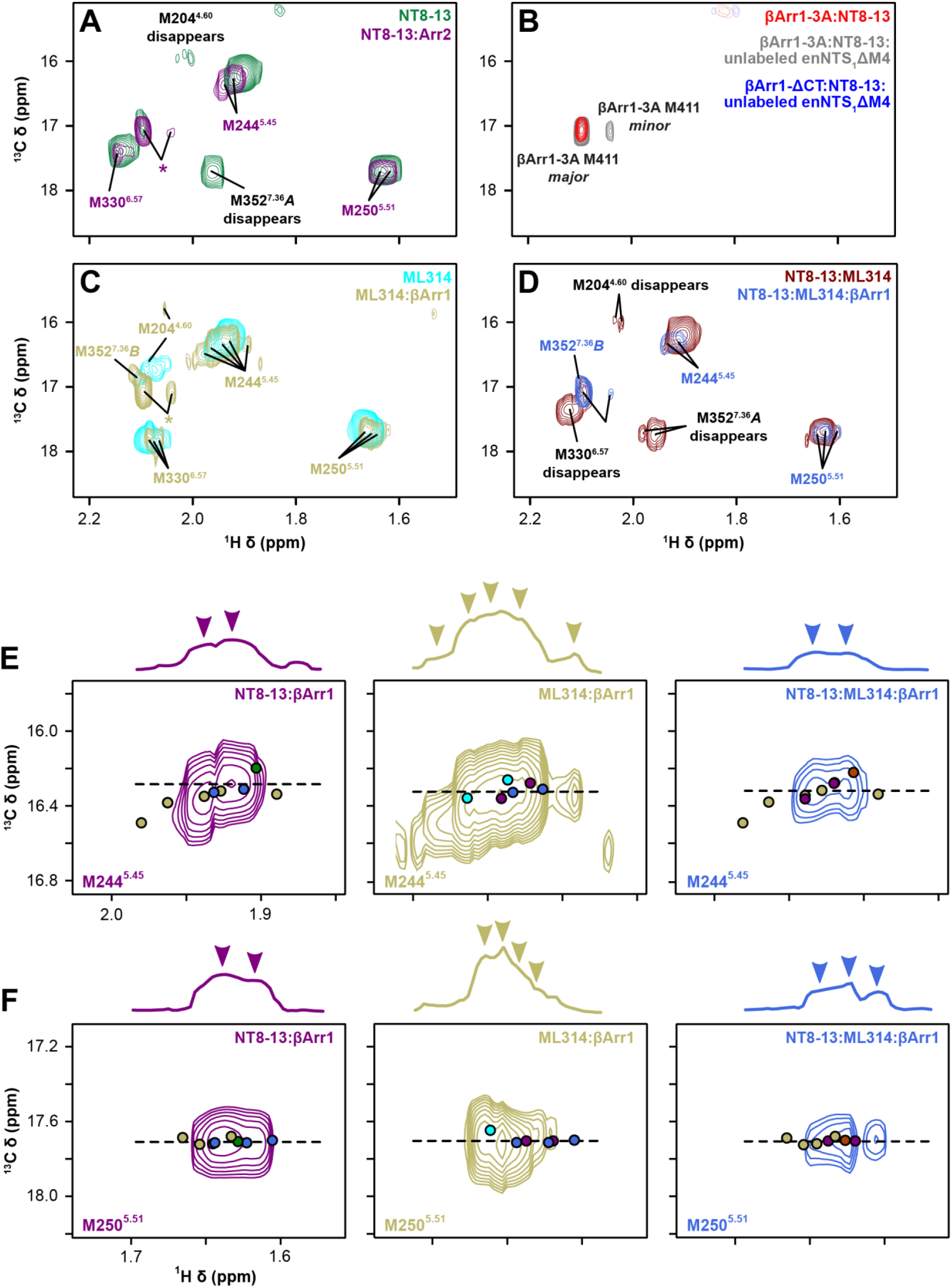
βArr1 stabilizes pre-existing states in the presence of agonist and/or BAM. A) Overlay of NT8-13:enNTS_1_ΔM4 (magenta) and NT8-13:enNTS_1_ΔM4:βArr1-3A (cyan) ^1^H-^13^C HMQC spectra; asterisks indicate natural abundance βArr1-3A M411 peaks. B) Comparison of 87 μM βArr1-3A (red), 165 μM βArr1-3A + 55 μM unlabelled-enNTS_1_ΔM4 (grey), and 150 μM βArr1-ΔCT + 65 μM unlabelled-enNTS_1_ΔM4 (blue) ^1^H-^13^C HMQC spectra. All spectra were collected in the presence of NT8-13 and PIP2 in DDM micelles. The two βArr1 resonances were both assigned to M411 because of their absence in the βArr1-ΔCT spectrum (blue) where the protein was truncated at N382. The minor βArr1 M411 resonance is only visible in the presence of enNTS_1_ΔM4 (grey), suggesting that it reflects a receptor-bound conformation. Overlays of ML314 (C) and NT8-13:ML314 (D) bound enNTS_1_ΔM4 ^1^H-^13^C HMQC spectra with and without 2.3x molar equivalents βArr1-3A. Peaks marked with an asterisk represent natural abundance βArr1-3A M411. Extracted spectral region of (D) M244^5.45^ and (E) M250^5.51^ from NT8-13:enNTS_1_ΔM4:βArr1-3A (purple), ML314:enNTS_1_ΔM4:βArr1-3A (tan), and NT8-13:ML314:enNTS_1_ΔM4:βArr1-3A (royal blue) ^1^H-^13^C HMQC spectra. One dimensional ^1^H cross-sectional slices (corresponding to dotted line) shown on top. Dots denote the residue’s chemical shift position in spectra of the corresponding colour with additional dots shown for ligand-only spectra (NT8-13:enNTS_1_ΔM4, forest green; ML314:enNTS_1_ΔM4, cyan; NT8-13:ML314:enNTS_1_ΔM4, maroon). All spectra were recorded at 600 MHz with receptor concentrations of 66 μM.

βArr1-3A leads to exchange broadening of M204^4.60^, M352^7.36^ state A and presumably M352^7.36^ state B, although the latter is overlapped with the major βArr1-3A M411 resonance (Figure 2A; asterisks). To better understand the chemical exchange kinetics, we performed a titration series with increasing βArr1-3A concentrations added to separate, otherwise identical, NT8-13:enNTS_1_ΔM4 samples (Figure S3). The intensity of every resonance decreased in a concentration-dependent manner reflecting the ternary complex’s longer rotational correlation time. The especially rapid broadening of M352^7.36^ indicates that arrestin back-coupling enhances μs-ms exchange kinetics at the periphery of the orthosteric pocket (Figure S3); this is consistent with the lack of density for the homologous hNTS_1_ M346^7.36^ sidechain in both NT8-13:hNTS_1_:βArr1 cryo-EM models^22,24^. The three methionines nearest to the transducer interface (M204^4.60^, M244^5.45^, and M250^5.51^) all split into at least two distinct conformational states exchanging on the ms-s timescale (Figure S3G-I). The major M244^5.45^ peak settled at a chemical shift position linearly between the apo- and NT8-13 bound states (Figure S3H) with a second peak produced at a similar chemical shift observed for the apo- and ML314 bound states (Figure 1B,D). βArr1-3A splits M250^5.51^ into two peaks centered at the NT8-13-bound chemical shift (Figures 1C, 2A, Fig S3I). Taken together, this indicates βArr1-3A is modulating the exchange kinetics of pre-existing agonist-bound conformations of the PIF motif – consistent with the proposed β2-adrenergic receptor activation mechanism^48^.

### BAM potentiates the exchange dynamics of the βArr-ternary complex

To test if exchange kinetics of the receptor’s conformational ensemble plays a general role in arrestin activation, we investigated ML314 ternary and ML314:NT8-13 quaternary complexes. The ML314:enNTS_1_ΔM4:βArr1-3A complex spectrum was qualitatively quite similar to NT8-13:enNTS_1_ΔM4:βArr1-3A with enhanced μs-ms exchange peripheral to the orthosteric pocket and slower ms-s motions near to the transducer interface (Figure 2A,C). Yet, there were several key differences. M352^7.36^ state B remained visible at 2.3 molar equivalents βArr1-3A and even increased in intensity relative to ML314:enNTS_1_ΔM4 (Figure 2C); thus βArr1-3A stabilizes the ML314 bound enNTS_1_ΔM4 conformer, which we hypothesize reflects a detached N-terminus and tightly packed TM1/TM2/TM7 interface^37^. M244^5.45^ splits into at least five peaks including enNTS_1_ΔM4 substates populated in the presence of ML314 and NT8-13:βArr1-3A (Figures 2A,E) as well as the SR142948A antagonist^37^. ML314 maintains M250^5.51^ in the furthest downfield position of any ligand with βArr1-3A perturbing it even further and simultaneously splitting it into an ensemble of at least three substates (Figures 2C,F). As βArr1-3A also pushes M250^5.51^ downfield relative to NT8-13:enNTS_1_ΔM4, we hypothesize this chemical environment signifies a transducer-competent conformer (Figure 2A). The simultaneous addition of ML314 and NT8-13 to enNTS_1_ΔM4:βArr1-3A collapses both M244^5.45^ and M250^5.51^ to a subset of resonances that more generally reflect a concatenation of the ML314:βArr1-3A and NT8-13:βArr1-3A ternary complexes (Figure 2).

### NTS_1:_Gα_iq_ conformational and kinetic ensemble is distinct from NTS_1:_βArr1-3A

NTS_1_ can uniquely couple to all major Gα protein subtypes (Gα_q/11_, Gα_i/o_, Gα_s_, and Gα_12/1349_) with the strongest preference towards G_q_ activation (Besserer-Offroy et al., 2017). This reflects a higher affinity and nucleotide exchange rate for Gα_q_, at least compared to Gα_i_, primarily driven by the six C-terminal residues of helix 5^50^. Since Gα_q_ is inherently unstable^51,52^, we took advantage of the Gα_iq_ chimera originally used to demonstrate that coupling specificity can largely be reduced to the G protein C-terminus^50^. NT8-13:enNTS_1_ΔM4:Gα_iq_ complex formation was supported by Gα_iq_ concentration-dependent changes in 1D ^1^H and 2D ^1^H-^13^C HMQC spectra (Figures 3A,B and S4). Two additional resonances, originating from unlabeled Gα_iq_ protein, are observed in the 2D ^1^H-^13^C HMQC spectrum but cannot be assigned to specific residues (Figure S5A).

**Figure 3.**
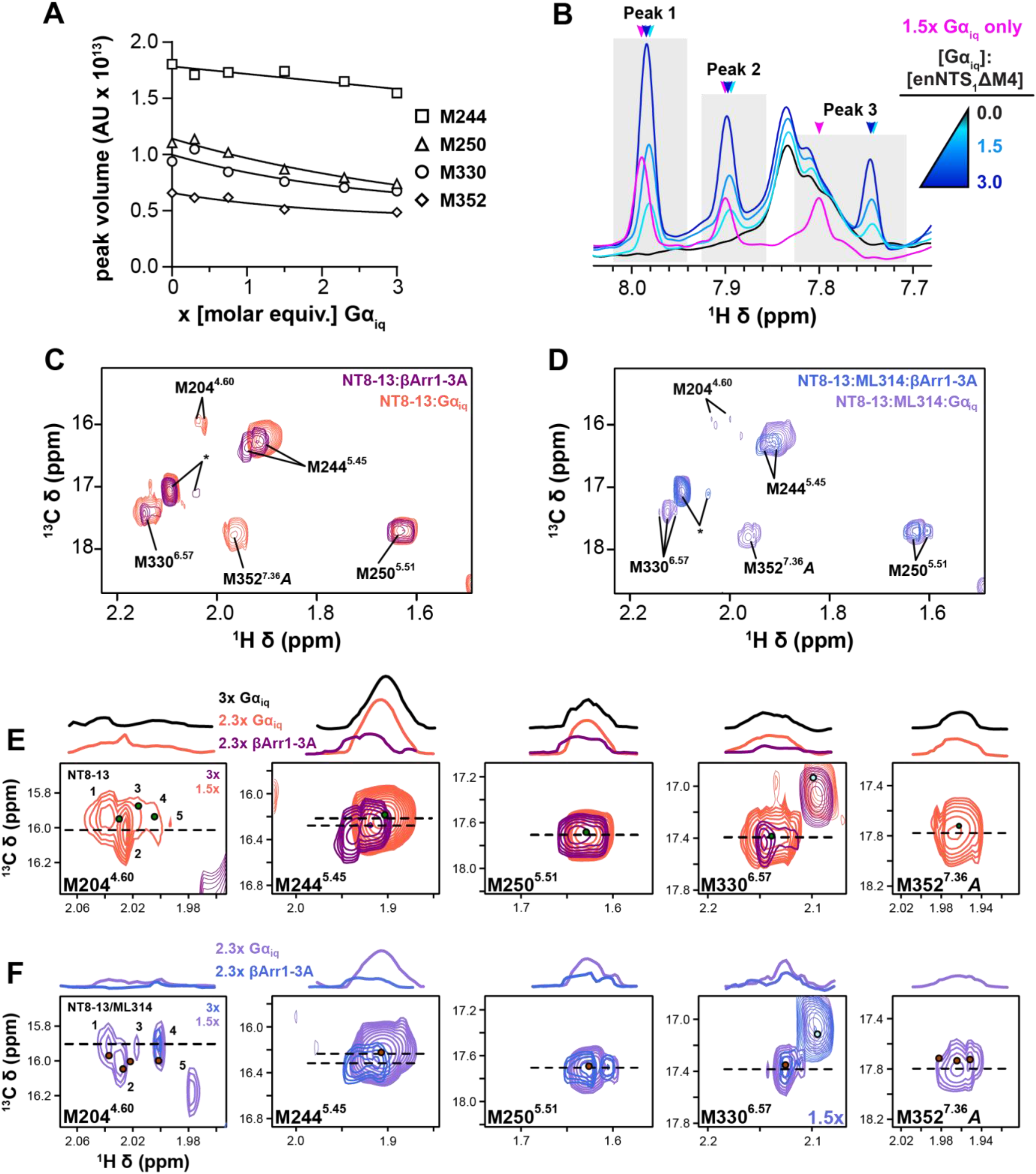
Comparison of Gα_iq_:enNTS_1_ΔM4 and βArr1-3A:enNTS_1_ΔM4 ternary complex NMR spectra. A) Integrated peak volumes of selected enNTS_1 13_C^ε^H_3_-methionine peaks at 0, 0.3x, 0.75x, 1.5x, 2.3x and 3x molar excess of Gα_iq_ indicate complex formation. B) ^1^H 1D spectra of three unassigned Gα_iq_ resonances exhibiting enNTS_1_ΔM4-dependent chemical shift perturbations. C) Overlay of NT8-13:enNTS_1_ΔM4:βArr1-3A (purple) and NT8-13:enNTS_1_ΔM4:Gα_iq_ (tomato) ^1^H-^13^C HMQC spectra. Peaks marked with an asterisk represent natural abundance βArr1-3A and Gα_iq_ resonances (Figures S12A and S16D). D) Overlay of NT8-13:ML314:enNTS_1_ΔM4:βArr1-3A (royal blue) and NT8-13:ML314:enNTS_1_ΔM4:Gα_iq_ (medium purple) ^1^H-^13^C HMQC spectra. Peaks marked with an asterisk represent natural abundance βArr1-3A and Gα_iq_ resonances. Extracted spectra regions of enNTS_1_ΔM4:Gα_iq_ and enNTS_1_ΔM4:βArr1-3A methionine resonances in the presence of E) NT8-13 and F) NT8-13:ML314. One dimensional ^1^H cross-sectional slices (corresponding to dotted line) shown on top. Dots denote the residue’s chemical shift position in spectra of the corresponding colour with additional dots shown for ligand-only spectra (NT8-13:enNTS_1_ΔM4, forest green; NT8-13:ML314:enNTS_1_ΔM4, maroon). Gα_iq_ containing spectra were recorded at 600 MHz with receptor concentrations of 64 μM and βArr1-3A containing spectra with receptor concentration of 66 μM.

A comparison of the Gα_iq_ and βArr1-3A titration series is particularly revealing (Figures 3C-F, S3 and S4). M250^5.51^ and M330^6.57^ concomitantly split into multiple, overlapping resonances in both ternary complexes, but βArr1-3A qualitatively promotes these changes at a slightly lower relative concentration. It is reasonable to anticipate a subset of similarly behaving resonances considering the highly conserved receptor architecture observed across all transducer complex structures and presumably partially-overlapped allosteric coupling networks^22-25^. There are three striking differences between Gα_iq_ and βArr1-3A ternary complex spectra. First, M244^5.45^ remains a single unperturbed resonance in the presence of up to 3x molar excess of Gα_iq_. We do not observe any changes, apart from a subtle intensity reduction, suggesting that i) NT8-13 alone induces a fully-active, G protein-competent M244^5.45^ conformation; ii) the isolated Gα_iq_ subunit is insufficient to stabilize the fully-active state; or iii) that M244^5.45^ does not play a role in Gα_iq_ coupling. Secondly, both transducers split M352^7.36^ state A into at least two resonances but βArr1-3A leads to exchange broadening at lower concentrations (Figures 3C-F, S3K, and S4K). Finally, Gα_iq_ stabilizes three M204^4.60^ resonances while βArr1-3A broadens all peaks before selecting a single state at 3x molar equivalents (Figures 3C-F, S3G, and S4G). A detailed comparison of M204^4.60^ is challenging due to the overall weak intensities and similar resonance patterns observed at sub-stoichiometric transducer concentrations (Figures 3C-F, S3G, and S4G). Yet, taken together, these chemical shift and intensity changes suggest that Gα_iq_ and βArr1-3A differentially modulate the kinetics of enNTS_1_ΔM4 conformational ensembles near the connector region and orthosteric binding pocket.

### BAM stabilizes a distinct Gα_iq_ quaternary complex

We collected a ML314:NT8-13:enNTS_1_ΔM4:Gα_iq_ spectrum to determine the mechanism by which ML314 attenuates G protein activation. We observe differential effects across the receptor with residues adjacent to the orthosteric pocket appearing to adopt more βArr1-competent-like conformers and those near the transducer-binding interface continuing to populate Gα_iq_-competent conformers. For example, ML314 reduces the M352^7.36^ state A intensity as observed in βArr1 ternary and quaternary complexes (Figures 3D-F, S3 and S5). At the bottom of the orthosteric pocket, ML314 again pushes M204^4.60^ along a linear trajectory towards a state that is only observed in βArr1-3A ternary and quaternary complexes, we hypothesize this may affect the hydrogen-bond network^37^ that governs receptor activation (Figure S3G). M244^5.45^ shows no substantial difference, apart from a subtle reduction in peak intensity, between NT8-13:enNTS_1_ΔM4, ML314:NT8-13:enNTS_1_ΔM4 and NT8-13:enNTS_1_ΔM4:Gα_iq_ (Figures S5). Whereas residue M250^5.51^, the closest probe to the transducer interface, begins to reflect a concatenation of Gα_iq_ and βArr1-competent states. Perhaps the most dramatic change is observed in the M330^6.57^ multiplet pattern. While titration of either transducer initially splits M330^6.57^ in the ^1^H dimension, βArr1 ultimately stabilizes the downfield resonance (Figures S3 and S4). However, when ML314 is present, regardless of transducer and/or NT8-13 combination, the upfield peak is stabilized which we hypothesize indicates nearby ML314 binding rather than receptor pharmacology (Figures 2C and 3F).

## DISCUSSION

Over the last two decades, extensive crystallographic and cryo-electron microscopy studies have laid a structural foundation for GPCR activation in terms of inactive, intermediate and active-state models. Sophisticated spectroscopic and computational studies expanded the conformational landscape to include high energy intermediate states from the fs-ns fluctuation of bond angles and side chain rotamers^53^ to the ns-μs toggling of microswitches^34,48,54,55^, the μs-ms conformational exchange of secondary structure^33,56^, and the ms-s activation of transducers^56,57^. Site-selective NMR labeling strategies, which are sensitive to molecular motions over the picosecond to second timescale, have proven especially powerful for describing how ligands and transducers remodel the GPCR conformational landscape^26,58^. Here, we employed endogenous ^13^C^ε^H_3_-methionine probes located around the extracellular vestibule and near the connector region to expand our understanding of how ligands, allosteric modulators and transducers regulate NTS_1_ motions.

X-ray crystallography and MD studies suggest that ligand binding is communicated to the transducer interface through correlated motions near the connector or transmission region^48,59^. We observe modest pairwise peak intensity correlations between these regions that are dramatically strengthened upon addition of PIP2 (Figure 4A). The spatial separations between the intracellular PIP2 binding pocket, the connector region, and orthosteric ligand provide strong evidence for a long-range allosteric link. We attribute these differential peak intensities to motions on the microsecond-millisecond timescale (i.e. exchange broadening). Although it is also possible that the ps-ns motions suggested by DFT analysis^37,44^ could also result in increased peak intensities through a longitudinal (T_1_) relaxation mechanism^60^, we hypothesize that the SOFAST-HMQC experiments employed here greatly reduce that possibility^61^. Nonetheless, future experiments will be required to quantitate the T_1_, T_2_, and generalized order parameters for each methionine methyl group.

**Figure 4.**
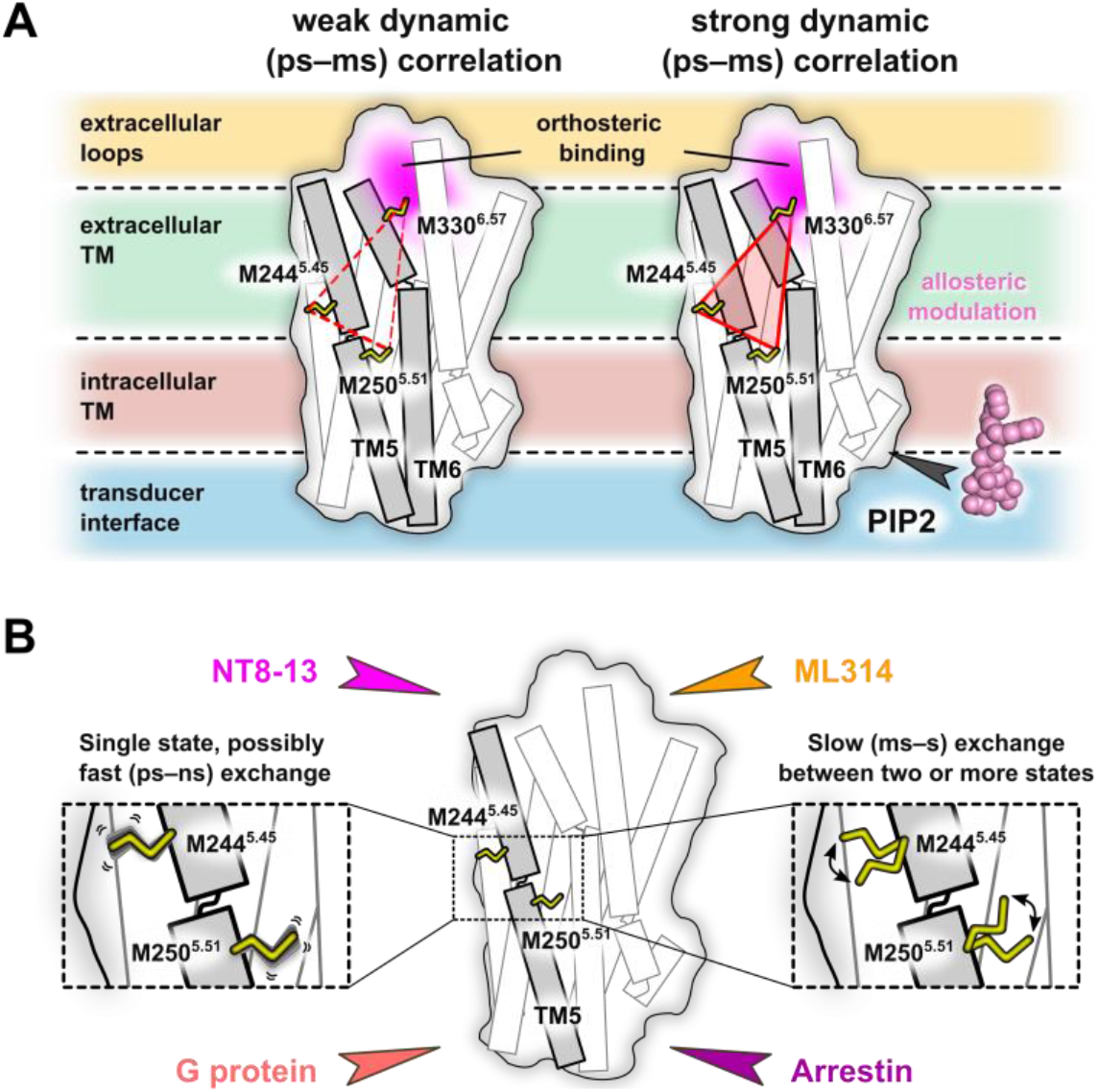
Ligands and transducer remodel the enNTS_1_ kinetic and/or thermodynamic ensemble. A) The weak pair-wise correlation of peak intensities near the orthosteric pocket and connector region are strengthened by PIP2 to reveal several long-range allosteric communication pipelines. B) β-arrestin-1 association slows the timescale of M244^4.45^ and M250^5.51^ conformational exchange whereas G protein coupling has little to no effect. Our NMR spectra qualitatively suggest the pre-existence of transducer-competent conformations in the agonist-bound state and that ML314, a β-arrestin biased allosteric modulator (BAM), fine-tunes exchange between those states.

Linking our observation to a specific enNTS_1_ΔM4:PIP2 interface is challenging. A hNTS_1_:βArr1ΔCT cryo-EM structure^22^ and a native mass spectrometry (MS) approach^36^ both indicate PIP2 binds at the TM1/2/4 groove; however, the same MS study proposes an additional interface formed by TM1/7^36^. Binding is dominated by electrostatic interactions between the polyanionic phosphorylated inositol head group of PIP2 and basic residues (Arg and Lys) of the receptor^22,36,62,63^. In the case of enNTS_1_, directed evolution resulted in three mutations (N262^5.63^R, K263^5.64^R and H305^6.32^R) at the intracellular tips of TMs 5 and 6, which could elevate the relevance of this second site^64,65^. NT8-13:ML314:enNTS_1_ΔM4 was the only ligand complex to exhibit substantial chemical shift changes upon addition of PIP2 (Fig. 1E). Perhaps this reflects counteracting allosteric networks, or perhaps even direct binding competition, of a balanced (PIP2) and biased (ML314) ligand in the absence of transducer.

Recent cryo-EM structures of hNTS_1_:βArr1 and hNTS_1_:Gα_i_βγ ternary complexes possess very high receptor structural similarity, which raises questions as to the role of conformational dynamics in functional selectivity. Although technically challenging, other studies have also begun to integrate kinetic information for a more complete description of the conformational landscape^66-68^. Both M244^4.45^ and, to a lesser extent, M250^5.51^ resonances split in response to βArr1 but not Gα_iq_ (Figures 3F and 4B). This peak pattern is indicative of a chemical exchange process on the slow NMR timescale suggesting a reasonably high activation barrier between substates. Interestingly, the βArr-biased allosteric modulator ML314 alone (or in combination with NT8-13) also induces splitting of those resonances, suggesting pre-selection of βArr-binding-competent states and offering a mechanistic basis for transducer-bias. Although speculative, we hypothesize that peak splitting in the ^1^H dimension may originate from local fluctuations of one or several aromatic rings^37,44^. Finally, it is interesting to note the continued presence of Gα_iq_ competent chemical shifts in the ML314:NT8-13:enNTS_1_ΔM4:Gα_iq_ spectrum (Fig. 3C-F). Similar studies using enNTS_1 19_F-labeled at the cytoplasmic tip of TM6 (Q301C^6.28^)^32^ reveal that a peptide corresponding to the G_q_ α5-helix binds ML314:enNTS_1_ΔM4 complexes with high-affinity by stabilizing unique TM6 conformers (J.J.Z., personal communication). Taken together, we hypothesize that ML314, and its successor SBI-553^69,70^, may bias NTS1 pharmacology by stabilizing signaling incompetent G protein conformations, such as a non-canonical Gα subunit^23,25^ and/or α5-helix pose^71,72^.

### Limitations of this study

Our model system employed several strategies to minimize the challenges to solution NMR studies of GPCR complexes such as multiple simultaneous binding partners, heterogenous phosphorylation patterns, high molecular weights, and inherent instability. We employed the pre-activated βArr-3A variant to control for uncertainty related to the number and position of phosphorylated residues in the receptor C-terminus. To minimize line-broadening side effects of slowly tumbling systems, we employed DDM detergent micelles and focused on only the Gα subunit that comprises nearly the entire complex interface. The Gα_iq_ chimera provides a robust scaffold to explore an otherwise unstable cognate receptor:G protein pair^51,52^. Thermostabilized enNTS_1_ permits extended data acquisition times that would otherwise be impossible; it binds ligands with similar affinity to rNTS_173_, and couples directly to G protein and β-arrestin, although with reduced affinity. Future studies will explore reversion of thermostabilizing mutations to further recover wildtype signaling capabilities. Nonetheless, loss-of-function mutations are more common than gain-of-function phenotypes suggesting that the molecular mechanisms of enNTS_1_/transducer coupling represent native allosteric pipelines. A long-term aim is to couple quantitative sidechain motions with all atom molecular dynamics to map allosteric connection pathways. Accurate quantitation requires highly deuterated systems^27,74^ that can be achieved by elegant means^75-77^, but are most easily afforded by *E. coli* expression systems.

## ACKNOWLEDGEMENTS

We are grateful to Prof. Aashish Manglik (UCSF) for providing the βArr1 construct used in this study. We would like to thank Prof. Andrew L. Lee (UNC) for fruitful discussions. The 14.1 T spectrometer at Indiana University used in this study was generously supported by the Indiana University Fund. The project was funded by: KAKENHI 21H04791 (AI), 21H051130 (AI), and JPJSBP120213501 (AI) from Japan Society for the Promotion of Science (JSPS); LEAP JP20gm0010004 (AI) and BINDS JP20am0101095 (AI) from the Japan Agency for Medical Research and Development (AMED); FOREST Program JPMJFR215T (AI) and JST Moonshot Research and Development Program JPMJMS2023 (AI) from Japan Science and Technology Agency (JST); Daiichi Sankyo Foundation of Life Science (AI); Takeda Science Foundation (AI); Ono Medical Research Foundation (AI); Uehara Memorial Foundation (AI); Agencia Estatal de Investigación, Spain grant PID2019-104914RB-I00 (JCP, MP); Australian National Health and Medical Research Council (NHMRC) grants 1081844 and 1141034 (RADB, DJS and PRG); Indiana Precision Health Initiative (JJZ); and National Institutes of Health (NIH) grants R00GM115814 (JJZ) and R35GM143054 (JJZ).

## METHODS

### *E. coli* expression and purification of enNTS_1_ variants

The protocols for ^13^C^ε^H_3_-methionine labelled expression of enNTS_1_ variants used for all NMR experiments as well as unlabeled expression in rich media used for thermostability and binding assays have previously been described in depth^73^. Expressions were usually carried out in batches of 3 L or 4 L and cell pellets were kept frozen at -80°C until further use. enNTS_1_ΔM4 (M204^4.60^/M208^4.64^/M244^5.45^/M250^5.51^/M330^6.57^/M352^7.36^) was purified as previously described^37^. Elutions from the initial IMAC capture step were directly cleaved with His-tagged HRV 3C protease (produced in-house) prior to concentrating using an Amicon 30 kDa MWCO concentrator (Millipore) and dilution with ion exchange chromatography (IEX) loading buffer (20 mM HEPES pH 8.0, 10% Glycerol, 0.02% DDM) to obtain a combined NaCl/Imidazole/Na_2_SO_4_ concentration of less than 50 mM. The cleaved receptor solution was then loaded onto a 5 mL HiTrap SP HP column (GE Healthcare) using an Akta Start system (GE Healthcare) and washed with the same buffer until the signal remained stable. The column was then washed with four column volumes of IEX wash buffer (20 mM HEPES pH 7.4, 10% Glycerol, 63 mM NaCl, 0.02% DDM) after which a 1 mL Ni-NTA HisTrap column (GE Healthcare) was inserted after the HiTrap SP HP column and the system was washed with another 10 mL of IEX wash buffer containing 10 mM Imidazole. The cleaved receptor was the eluted with IEX elution buffer (20 mM HEPES pH 7.4, 10% Glycerol, 1 M NaCl, 0.03% DDM, 20 mM Imidazole) and the receptor containing fractions concentrated to approx. 400 uL for injection onto a S200 Increase SEC column (GE Healthcare) using a 500 μL loop and an Akta Pure System (GE Healthcare). The receptor containing fractions from SEC purification using SEC buffer (50 mM Potassium phosphate pH 7.4, 100 mM NaCl, 0.02% DDM) were then concentrated and buffer exchanged (for NMR experiments) using NMR buffer (50 mM Potassium phosphate pH 7.4, 100 mM NaCl in 100% D_2_O) to reduce the residual H_2_O concentration to <1%. Receptor samples were then aliquoted and stored at -80 °C until further use. The modified purification protocol comprising the IEX step was found to yield a similar if not higher receptor purity compared to the original protocol containing a reverse IMAC step (Bumbak et al., 2019) as judged by SDS-Page. enNTS_1_ΔM4 used in NMR experiments retains a C-terminal Avi-tag (which was used for capture in ligand-binding and thermostability assays) and the amino acid sequence is: GPGSTSESDTAGPNSDLDVNTDIYSKVLVTAIYLALFVVGTVGNGVTLFTLARKKSLQSLQSRVDYYLGSLALSSLLILLFALPVDVYNFIWVHHPWAFGDAGCKGYYFLREACTYATALNVVSLSVERYLAICHPFKAKTLLSRSRTKKFISAIWLASALLSLPMLFTMGLQNLSGDGTHPGGLVCTPIVDTATLRVVIQLNTFMSFLFPMLVASILNTVIARRLTVLVHQAAEQARVSTVGTHNGLEHSTFNVTIEPGRVQALRRGVLVLRAVVIAFVVCWLPYHVRRLMFVYISDEQWTTALFDFYHYFYMLSNALVYVSAAINPILYNLVSANFRQVFLSTLASLSPGWRHRRKKRPTFSRKPNSVSSNHAFSTASGLNDIFEAQKIEWHEGSGLEVLFQ

### Expression and purification of βArr1-3A

The pET15 expression plasmid harboring the hβArr1-3A gene was a kind gift from Ashish Manglik. In this plasmid the hβArr1-3A sequence was modified previously by mutating I386A, V387A, F388A (termed 3A mutant)^78^ and by mutating 6 cysteine residues to other amino acid types (i.e. C59V, C125S, C140L, C242V, C251V and C269S). hβArr1-3A gene was preceded by a 6x His tag, HRV 3C protease cleavage site and Protein C tag. This sequence was modified by inserting an additional HRV 3C protease cleavage site between the Protein C sequence and the hβArr1-3A gene to allow complete removal of N-terminal tags. 5 mL of a LB day pre-culture containing 100 mg/L carbenicillin and 1% (w/v) glucose were inoculated with a single colony of *E. coli* BL21(DE3) cells (Lucigen, Middleton, WI) freshly transformed with the βArr1-3A expression plasmid. After 9 h (37 °C, 225 rpm) 10 μL of LB pre-culture were added to 50 mL of a Teriffic Broth (TB) pre-culture containing 100 mg/L carbenicillin and 1% (w/v) glucose, and incubated overnight at (30 °C, 225 rpm). The next morning the 50 mL of TB pre-culture were added to shaker flasks containing 950 mL of the same medium and incubated (37 °C, 225 rpm) to reach an OD600 of 0.6 at which point the temperature was reduced to 20 °C and the culture was incubated further until an OD600 of 1.0 was reached. The flasks were then cooled on ice for 5 min prior to induction with 0.4 mM isopropyl β-D-1-thiogalactopyranoside (IPTG). Protein expression was carried out at 20 °C and 225 rpm for 19h. The cells were harvested by centrifugation (5000 rcf, 4 °C, 15 min) and the combined pellets resuspended with wash buffer (25 mM HEPES, 100 mM NaCl, pH 8) and the washed cells were then pelleted by centrifugation (3000 rcf, 4 °C, 15 min) and stored at −80 °C. Thawed cells were resuspended in solubilization buffer (20 mM HEPES pH 8, 500 mM NaCl, 2 mM MgCl2, 15% glycerol, 1 Roche EDTA free Protease inhibitor tablet, 0.4 mM PMSF, 1 mg/mL Lysozyme, 1 uL/mL DNAse) and left stirring at 4 °C for 30 min prior to sonication on ice. Cell debris was removed by centrifugation (24000g, 4 °C, 45 min) and the supernatant filtered using a 45 μm syringe filter (Millipore). The filtrate was then incubated for 1 h rotating at 4 °C with 2 mL Ni-NTA resin (Thermo Fisher) per 1.5 L of expression culture. The resin was then washed with 15 mL wash buffer 1 (20 mM HEPES pH 8, 300 mM NaCl, 10% glycerol, 10 mM imidazole) followed by 12 mL wash buffer 2 (same as wash buffer 1 but 20 mM imidazole) and 12 mL wash buffer 3 (same as wash buffer 1 but 25 mM imidazole) per 1 mL of resin. βArr1-3A was eluted with approx. 10 mL of elution buffer (20 mM HEPES pH 7.5, 150 mM NaCl,10% glycerol, 200 mM imidazole) per 1 mL of resin. The 6x His-tag was removed via His-tagged HRV 3C protease (produced in-house) cleavage overnight rotating at 4 °C. The cleavage reaction was then concentrated using an Amicon 30 kDa MWCO centrifugal concentrator (Millipore) and diluted with 20 mM HEPES pH 7.5 to obtain a combined NaCl/Imidazole/Na_2_SO_4_ concentration of less than 50 mM. The solution was again filtered using a 45 μm syringe filter (Millipore) prior to loading onto a 5 mL HiTrap Q IEX column (GE Healthcare) equilibrated with 20 mM HEPES pH 7.5 followed by a wash step with the same buffer until a conductivity of 5 mS/cm was reached. The column was then further washed with IEX wash buffer (20 mM HEPES pH 7.5, 50 mM NaCl) until the A280 signal stabilized. The column was eluted using a 25 min gradient stretching from 50 mM to 500 mM NaCl. The βArr1-3A containing fractions were then pooled and concentrated to approx. 750 μL using an Amicon 30 kDa MWCO centrifugal concentrator (Millipore) prior to injection onto a HiLoad 16/600 S200pg SEC column (GE Healthcare) equilibrated with SEC buffer (20 mM HEPES pH 6.8, 150 mM NaCl) using a 1 mL loop. The βArr1-3A containing SEC fractions were pooled, concentrated to 296 μM and aliquots stored at -80°C until further use. The amino acid sequence of βArr1-3A used in NMR experiments is: GPSGDKGTRVFKKASPNGKLTVYLGKRDFVDHIDLVDPVDGVVLVDPEYLKERRVYVTLTVAFRYGREDLDVLGLTFRKDLFVANVQSFPPAPEDKKPLTRLQERLIKKLGEHAYPFTFEIPPNLPSSVTLQPGPEDTGKALGVDYEVKAFVAENLEEKIHKRNSVRLVIRKVQYAPERPGPQPTAETTRQFLMSDKPLHLEASLDKEIYYHGEPISVNVHVTNNTNKTVKKIKISVRQYADIVLFNTAQYKVPVAMEEADDTVAPSSTFSKVYTLTPFLANNREKRGLALDGKLKHEDTNLASSTLLREGANREILGIIVSYKVKVKLVVSRGGLLGDLASSDVAVELPFTLMHPKPKEEPPHREVPENETPVDTNLIELDTNDDDAAAEDFARQRLKGMKDDKEEEEDGTGSPQLNNR

### Expression and purification of Gα_iq_

The codon optimized gene for the Gα_iq_ chimera^50^ was purchased from GenScript (Piscataway, NJ) and subcloned into a pIQ expression vector with an open reading frame encoding an N-terminal 6xHis tag followed by a NNNNNNNNNNG linker, a MBP sequence and a HRV 3C protease cleavage site (LEVLFQGP). 50 mL of a LB day pre-culture containing 100 mg/L carbenicillin and 1% (w/v) glucose were inoculated with a single colony of *E. coli* BL21(DE3) cells (Lucigen, Middleton, WI) freshly transformed with the Gα_iq_ expression plasmid. After 9 h (37 °C, 225 rpm) 20 mL of LB pre-culture were centrifuged (3000 rcf, RT, 5 min) and the resuspended pellets were used to inoculate shaker flasks with 1 L of 2xYT medium containing 100 mg/L carbenicillin and 0.2% (w/v) glucose. The culture was incubated (37 °C, 225 rpm) to reach an OD600 of 0.7. The flasks were then cooled on ice for 5 min prior to induction with 1 mM IPTG. Protein expression was carried out at 25 °C and 225 rpm for 16 h. The cells were harvested by centrifugation (5000 rcf, 4 °C, 15 min) and the combined pellets resuspended with wash buffer (25 mM HEPES, 100mM NaCl, pH 8) and the washed cells were then pelleted by centrifugation (3000 rcf, 4 °C, 15 min) and stored at −80 °C. Gα_iq_ was purified following a protocol for purification of miniG proteins published previously (Carpenter and Tate, 2017). His-tagged HRV 3C protease (produced in-house) was used instead of TEV protease and a HiLoad 16/600 S200pg SEC column (GE Healthcare) was used instead of a HiLoad 26/600 S200 SEC column. The Gα_iq_ containing SEC fractions were pooled, concentrated to 721 μM and aliquots stored at -80°C until further use. The amino acid sequence of Gα_iq_ used in NMR experiments is: GPGSGCTLSAEDKAAVERSKMIDRNLREDGEKAAREVKLLLLGAGESGKSTIVKQMKIIHEAGYSEEECKQYKAVVYSNTIQSIIAIIRAMGRLKIDFGDSARADDARQLFVLAGAAEEGFMTAELAGVIKRLWKDSGVQACFNRSREYQLNDSAAYYLNDLDRIAQPNYIPTQQDVLRTRVKTTGIVETHFTFKDLHFKMFDVGGQRSERKKWIHCFEGVTAIIFCVALSDYDLVLAEDEEMNRMHESMKLFDSICNNKWFTDTSIILFLNKKDLFEEKIKKSPLTICYPEYAGSNTYEEAAAYIQCQFEDLNKRKDTKEIYTHFTCATDTKNVQFVFDAVTDVIIKNNLKEYNLV

### NMR spectroscopy

NMR spectra were collected on 600 MHz Bruker Avance Neo spectrometers equipped with a triple resonance cryoprobes. 2D ^1^H-^13^C SOFAST-HMQC spectra^61^ were recorded with 25% non-uniform sampling (NUS) at 298 K with a ^1^H spectral width of 12 ppm (1024 data points in t_2_) and a ^13^C spectral width of 25 ppm (128 data points in t_1_), relaxation delays of 450 ms, and 2048 scans per t1 data point resulting in acquisition times of 10 h per spectrum. A 2.25 ms PC9 120 degree ^1^H pulse^79^ was applied for excitation and a 1 ms r-SNOB shaped 180 degree ^1^H pulse^80^ was used for refocusing. The ^13^C carrier frequency was positioned at 17 ppm, and the ^1^H at 4.7 ppm, while band selective ^1^H pulses were centered at 1.8 ppm. 1D ^1^H spectra were recorded at 298 K with a spectral width of 13.7 ppm (2048 data points) and a relaxation delay of 1 s, and 128 scans. Samples were prepared to volumes of 160 μL in 3 mm tubes (Willmad), containing 20 μM DSS and 0.05% Na_2_N. Ligands were added to a final concentration of 500 μM. NT8-13 (5-10 mM) stock solutions were prepared in 100% D_2_O and ML314 (20 mM) in 100% DMSO-d6. PIP2 was added to a final concentration of 130 μM (approx. 2x molar equivalents of receptor). βArr1-3A and Gα_iq_ aliquots of 0.3-3x molar equivalents of receptor were buffer exchanged three times with NMR buffer (to >99%) prior to combining with the receptor. βArr1-3A containing samples were incubated for 1 h at room temperature prior to starting experiments. Gα_iq_ containing samples were supplemented with 2 mM MgCl_2_, 100 μM TCEP, and 10 μM GDP and incubated for 1 h at room temperature prior to adding 0.25 units of Apyrase (NEB) and incubation for a further 1 h at room temperature. All spectra were referenced against internal DSS, reconstructed with compressed sensing using qMDD^81^, and processed using NMRPipe^82^ where data were multiplied by cosinebells and zero-filled once in each dimension. Spectra were analyzed in Sparky (Goddard, T.D. and Kneller, D.G., University of California, San Francisco). All spectra from this study are reproduced together in Figure S6.

## Notes

### Competing Interest Statement

The authors have declared no competing interest.

