## Supplemental Information for "The dynamic nature of neurotensin receptor 1 (NTS_1_) allostery and signaling bias"

[1]Department of Molecular and Cellular Biochemistry, Indiana University, Bloomington, Indiana, USA 47405
[2]Graduate School of Pharmaceutical Sciences, Tohoku University, Sendai, Miyagi, Japan 980-8578
[3]Biomolecular NMR laboratory. Department of Inorganic and Organic Chemistry. Universitat de Barcelona (UB). 08028-Barcelona, Spain
[4]Department of Materials Science and Physical Chemistry & Institute of Theoretical and Computational Chemistry (IQTCUB). Universitat de Barcelona (UB). 08028-Barcelona, Spain
[5]Department of Biochemistry and Pharmacology, Bio21 Molecular Science and Biotechnology Institute, University of Melbourne, Parkville, Victoria 3010, Australia
[6]Department of Chemistry, Indiana University, Bloomington, Indiana 47405-7102, USA
[7]The Florey Institute of Neuroscience and Mental Health and Department of Biochemistry and Pharmacology, The University of Melbourne, Parkville, Victoria 3010, Australia
[8]Present address: ARC Centre for Cryo-electron Microscopy of Membrane Proteins and Drug Discovery Biology, Monash Institute of Pharmaceutical Sciences, Monash University, Parkville, Victoria 3052, Australia

**Keywords:** G protein-coupled receptors (GPCRs), NMR, allosteric modulator, conformational selection, Phosphatidylinositol-4,5-bisphosphate (PIP2), PIF motif, methionine

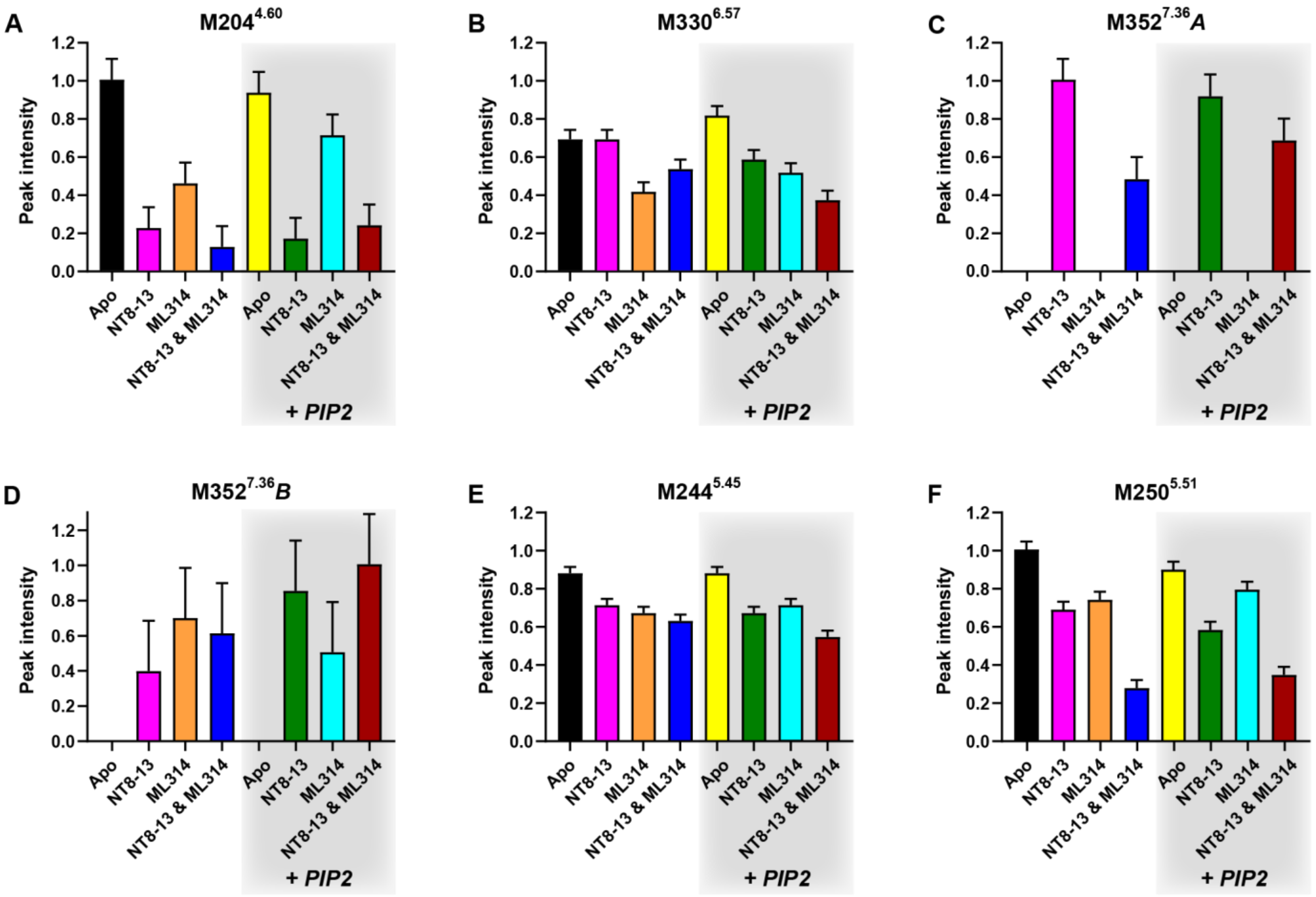


**Supplemental Figure S1. Effect of ligand and PIP2 combinations on enNTS$_1$ΔM4 $^{13}C^{\varepsilon}H_3$-methionine peak volumes.** Integrated peak volumes were determined by manual integration using Sparky**.** For each residue, peak volumes were normalized to the largest valued condition. Bar color correspond to the color scheme used for HMQC spectra. M204$^{4.60}$ intensities are the sum of all split resonances. Intensities for NT8-13 and NT8-13 & PIP2 conditions are the mean intensities from two independent samples. Error bars represent the standard deviation (SD) of peak volumes averaged for all 6 resonances in the NT8-13 and NT8-13 & PIP2 duplicate experiments.

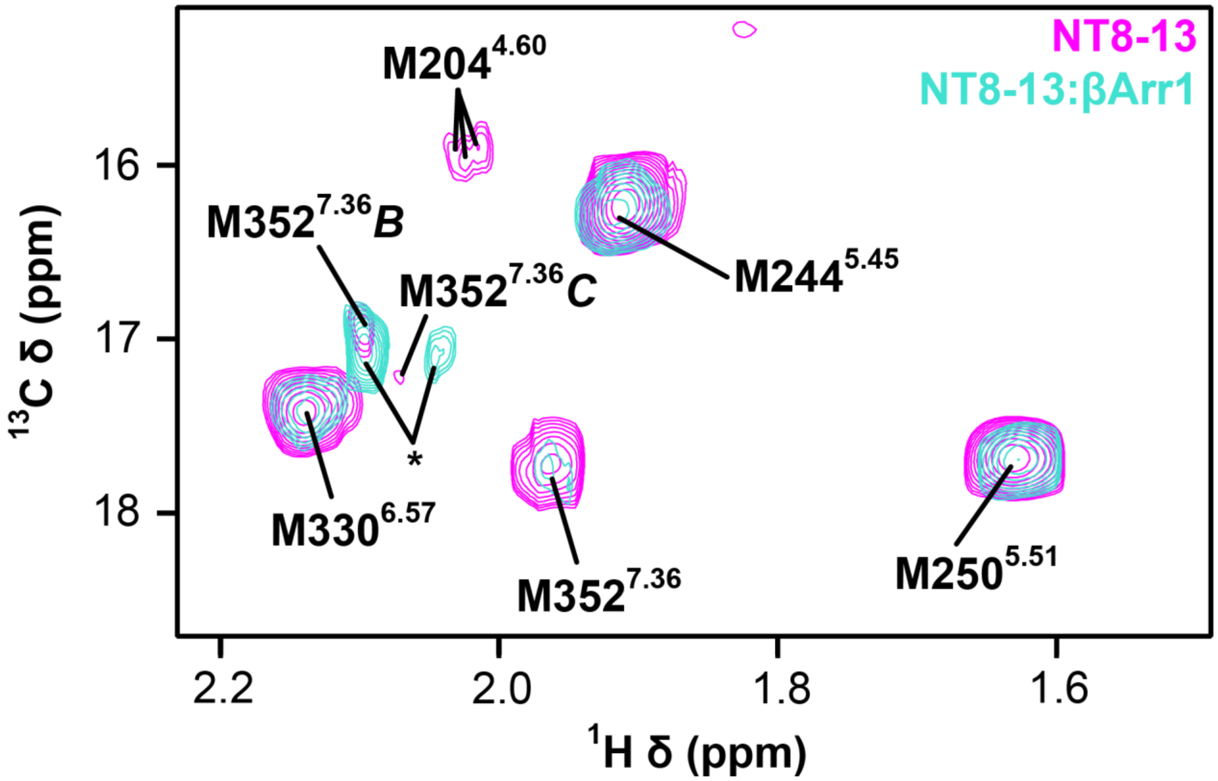


**Supplemental Figure S2. $^{1}$H-$^{13}$C HMQC spectra of [$^{13}C^{\varepsilon}H_3$-methionine]-enNTS$_1$ΔM4 binary and βArr1-3A ternary complexes in the absence of PIP2.** Overlay of NT8-13:enNTS$_1$ΔM4 (magenta) and NT8-13:enNTS$_1$ΔM4:βArr1-3A (cyan) $^{1}$H-$^{13}$C HMQC spectra; asterisks indicate natural abundance βArr1-3A M411 peaks. Both spectra were recorded at 600 MHz with enNTS$_1$ΔM4 concentrations of 66 μM and 2.3x molar excess βArr1-3A was used in the NT8-13:enNTS$_1$ΔM4:βArr1-3A experiment.

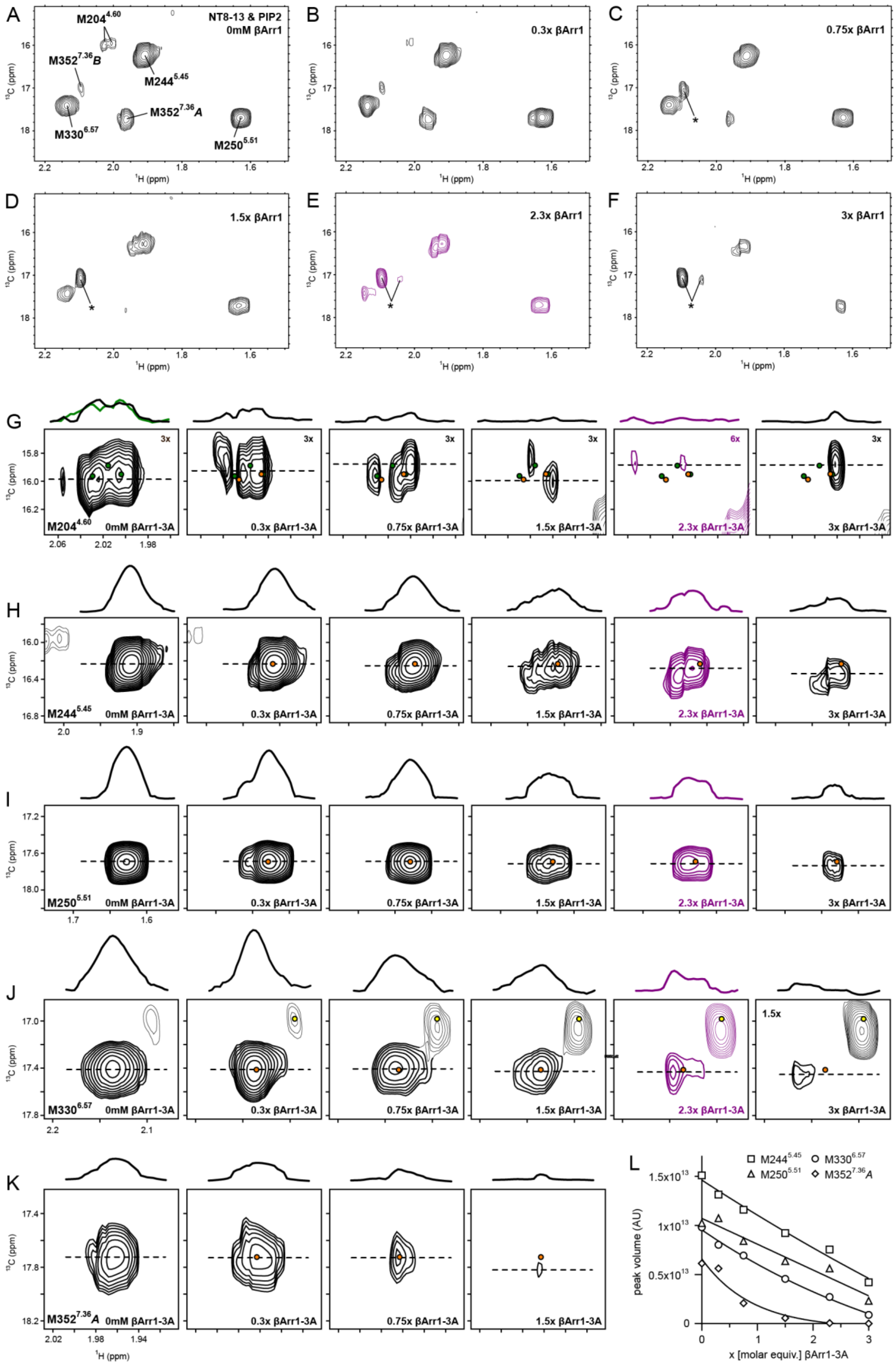


**Supplemental Figure S3. Titration of βArr1-3A into NT8-13:enNTS$_1$ΔM4:PIP2.** $^1$H-$^{13}$C HMQC spectra of enNTS$_1$ΔM4:NT8-13 in the presence of 0.0 (A), 0.3 (B), 0.75 (C), 1.5 (D), 2.3 (E), and 3.0 (F) molecular equivalents of βArr1-3A. Each spectrum was collected from separate, otherwise identical, NT8-13:enNTS$_1$ΔM4 samples. G-K) Extracted $^1$H-$^{13}$C HMQC spectral regions for individual methionine resonances. 1D $^1$H cross-sectional slices correspond to the dotted lines in the 2D spectra. All panels are plotted with identical contour levels unless otherwise indicated in the upper right corner (M204$^{4.60}$, 3x or 6x; M330$^{6.57}$, 1.5x). The resonances of other residues within the extracted region are drawn at 50% transparency.

Orange dots mark the positions of peaks in the absence βArr1-3A. Green dots (G) indicate peak positions observed in an independent experiment of enNTS$_1$ΔM4:NT8-13 without transducer (Supplemental Figure 4A and green panels of Supplemental Figures 4G-K). Yellow dots (J) mark the position of M352$^{7.36}$B which is obscured by the major βArr1-3A M411 peak. L) Absolute peak volumes of selected residues plotted against molecular equivalents of βArr1-3A suggest peak broadening as a function of complex formation. All spectra were recorded at 600 MHz, in 3 mm thin wall precision NMR tubes (Wilmad), with enNTS$_1$ΔM4 concentrations of 66 µM.

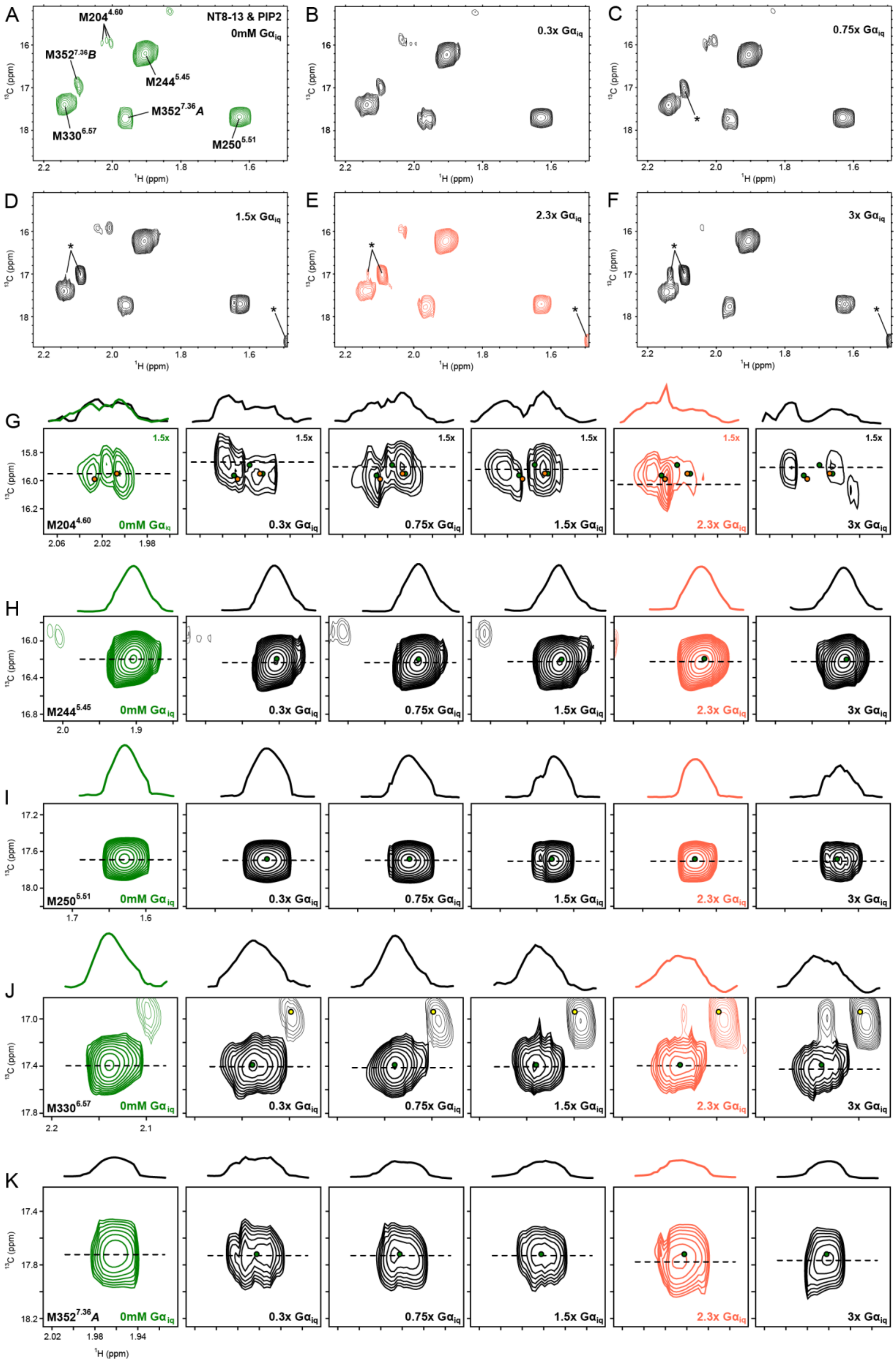


**Supplemental Figure S4. Titration of Gα$_{iq}$ into NT8-13:enNTS$_1$ΔM4.** $^{1}$H-$^{13}$C HMQC spectra of enNTS$_1$ΔM4:NT8-13 in the presence of 0.0 (A), 0.3 (B), 0.75 (C), 1.5 (D), 2.3 (E), and 3.0 (F) molecular equivalents of Gα$_{iq}$. Each spectrum was collected from separate, otherwise identical, NT8-13:enNTS$_1$ΔM4 samples. G-K) Extracted $^{1}$H-$^{13}$C HMQC spectral regions for individual methionine resonances. 1D $^{1}$H cross-sectional slices correspond to the dotted lines in the 2D spectra. All panels are plotted with identical contour levels unless otherwise indicated in the upper right corner (M204$^{4.60}$, 1.5x). The resonances of other residues within the extracted region are drawn at 50% transparency. Green dots mark the positions of peaks in the absence of $G\alpha_{iq}$. Orange dots (G) indicate peak positions observed in an independent experiment of NT8-13:enNTS$_1$ΔM4 without transducer (Supplemental Figure S3A and first panels of Supplemental Figures S3G-K). Yellow dots (J) mark the position of M352$^{7.36}$B which is obscured by the major natural abundance peak arising from $G\alpha_{iq}$. All spectra were recorded at 600 MHz, in 3 mm thin wall precision NMR tubes (Wilmad), with enNTS$_1$ΔM4 concentrations of 64 μM.

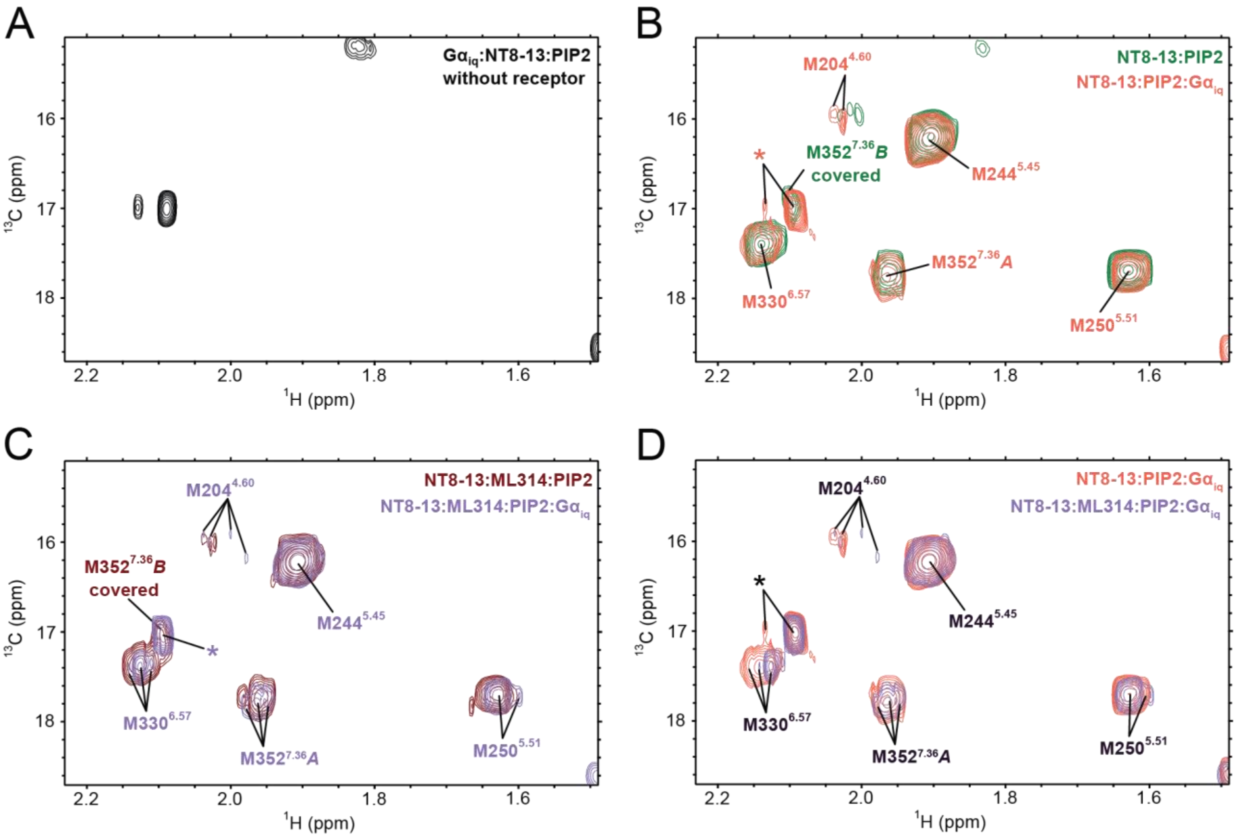


**Supplemental Figure S5. Comparison of Gα$_{iq}$ $^1$H-$^{13}$C HMQC spectra in the presence of various receptor and ligand combinations.** A) $^1$H-$^{13}$C HMQC spectrum of 99 μM Gα$_{iq}$, without receptor, suggest natural abundance $^{13}CH_3$-methionine peaks arising from Gα$_{iq}$. Overlays of NT8-13 (B) and NT8-13:ML314 (C) bound enNTS$_1$ΔM4 $^1$H-$^{13}$C HMQC spectra with and without Gα$_{iq}$. D) Comparison of NT8-13:enNTS$_1$ΔM4:PIP2:Gα$_{iq}$ (salmon) and NT8-13:ML314:enNTS$_1$ΔM4:PIP2:Gα$_{iq}$ (light purple) $^1$H-$^{13}$C HMQC spectra. Peaks marked with an asterisk represent natural abundance $^{13}CH_3$-methionine peaks arising from Gα$_{iq}$. All spectra were recorded at 600 MHz, in 3 mm thin wall precision NMR tubes (Wilmad), with enNTS$_1$ΔM4 concentrations of 64 μM.

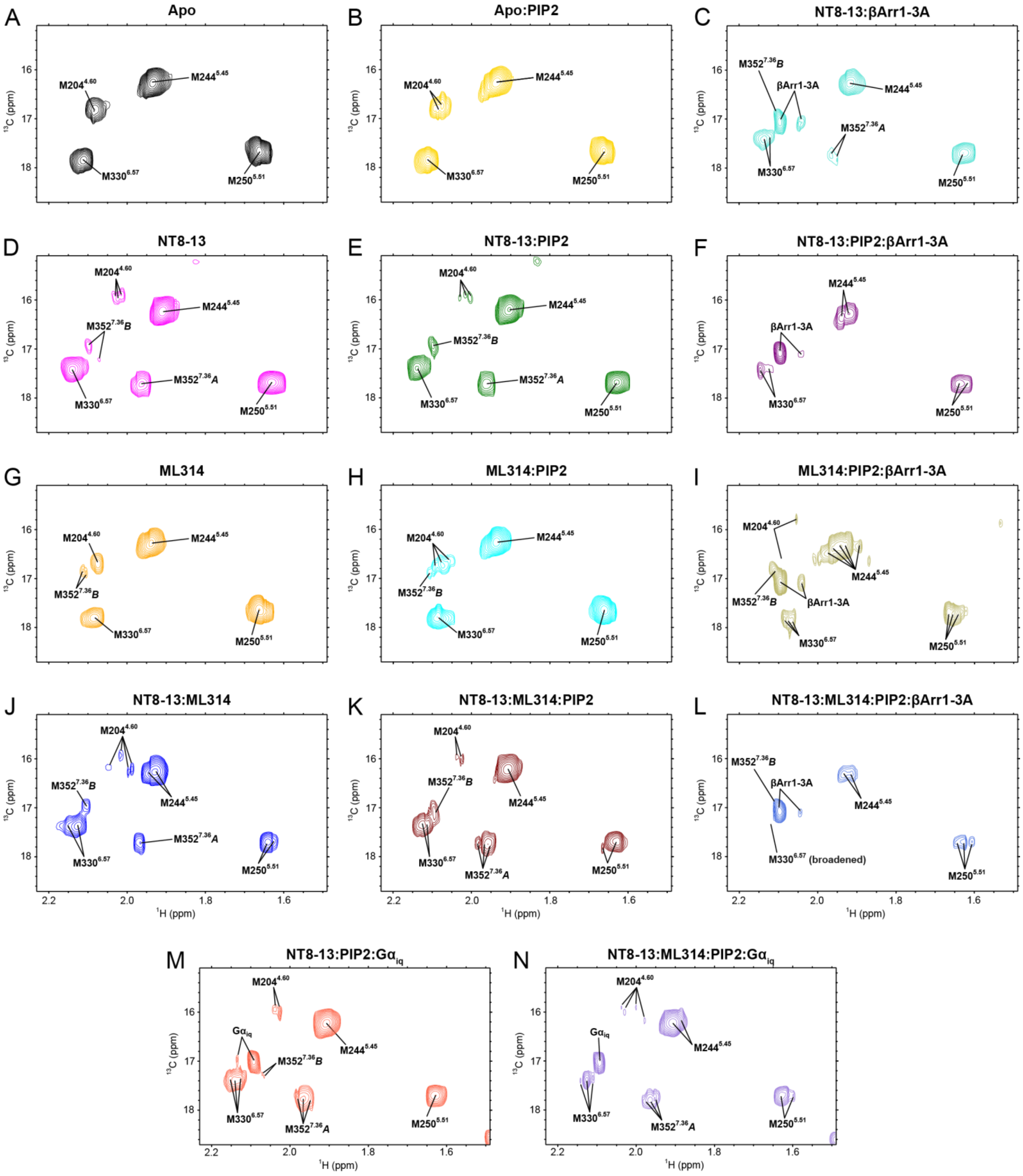


**Supplemental Figure S61. $^{1}H$-$^{13}C$ HMQC spectra of enNTS$_1$ΔM4 $^{13}C^{\varepsilon}H_3$-methionine for all ligand:transducer protein conditions.** Colour coding of individual spectra match those used throughout the manuscript. (F, I-O) Only spectra at 2.3x molar equivalents transducer are reproduced. All spectra were recorded at 600 MHz, in 3 mm thin wall precision NMR tubes (Wilmad), at enNTS$_1$ΔM4 concentrations of 66 μM (A-L) and 64 μM (M and N).
